# Challenges and promises in optimising a non-clinical protocol of intracerebroventricular human neural stem cell transplantation in ALS

**DOI:** 10.1101/2024.11.18.624143

**Authors:** Ivan Lombardi, Clelia Ferrero, Edvige Vulcano, Daniela Maria Rasà, Maurizio Gelati, Diego Pastor Campos, Rose Mary Carletti, Silvia de la Morena, Daniela Celeste Profico, Sabrina Longobardi, Elisa Lazzarino, Elisa Perciballi, Jessica Rosati, Salvador Martinez Perez, Alessandro Vercelli, Marina Boido, Daniela Ferrari

## Abstract

**Background and aims:** Neural stem cell (NSC) transplantation holds promising therapeutic potential for neurodegenerative disorders like amyotrophic lateral sclerosis (ALS). However, pre-clinical studies and early-phase clinical trials have faced challenges hindering the effective clinical translation of this approach. Crucial hurdles include the side-effects of prolonged immunosuppression, concerns regarding cell origin and transplantation dosage, identification of the most appropriate therapeutic window, and invasiveness of surgical procedures. Here, we show challenges and promises in optimizing a non-clinical protocol to assess safety and efficacy of human NSC (hNSC) intracerebroventricular (ICV) transplantation for ALS.

**Methods:** We evaluated the safety of administering up to 1×10^6^ hNSCs in immunodeficient mice and assessed their potential efficacy in reducing ALS hallmarks employing the SOD1^G93A^ mouse model. Both, transient (15 days) and prolonged immunosuppression regimens, at low (15 mg/kg) and high (30 mg/kg) doses, were tested along with two different cell dosages (3×10^5^ and 1×10^6^).

**Results:** Bilateral ICV injection of up to 1×10^6^ hNSCs proved to be safe, with no evidence of tumor formation. At 40 days post-transplantation, hNSCs induced a trend toward delaying motor decline and reducing spinal cord (SC) microgliosis when transplanted under prolonged high-dose (30 mg/kg) immunosuppression.

**Conclusions:** Our study suggests that: (i) a bilateral ICV transplantation of 1×10^6^ hNSCs is safe and non-tumorigenic in immunodeficient hosts; (ii) sustained high-dose immunosuppression is essential for ensuring cell survival in immunocompetent mice; and (iii) hNSC transplantation may provide therapeutic benefits in ALS by delaying motor decline and reducing microgliosis. This study also highlights persisting hurdles that need to be further addressed, such as the aggressive murine immune response to exogenous cells.

## Introduction

Amyotrophic lateral sclerosis (ALS), is a complex, multi-factorial and progressive neurodegenerative disorder^1^. It is characterized by the progressive degeneration of both primary motor neurons (MNs), in the motor cortex, and secondary motor neurons, in the brainstem and spinal cord (SC)^1^. Pathological hallmarks include MN degeneration and widespread neuroinflammation, both contributing to disease progression, culminating in extensive muscles denervation, paralysis and death mostly from respiratory failure within 2 - 5 years of diagnosis^1^. While the exact etiology remains elusive, 5 - 10% of ALS cases exhibit a familial, autosomal dominant inheritance pattern^2^ (fALS) linked to mutations in over 50 genes^3^, including *TARDBP*, *FUS* (∼5% of fALS); *SOD1* (∼20% of fALS) and *C9ORF72* (∼40% of fALS). Yet approximately 90% of ALS cases are sporadic (sALS), with unknown origin.

Several mechanisms have been proposed to contribute to ALS pathology. Among others, excitotoxicity, disruptions in cytoskeletal dynamics and axonal transport, altered RNA homeostasis, and non-neuronal cell function were demonstrated to contribute to neurodegeneration^4^. While proposed mechanisms shed light on ALS onset and progression, they also underscore the intrinsic challenges in developing effective therapies. Currently, treatment options are limited to symptomatic relief, with only few drugs offering modest survival benefits^5^.

Significant efforts have been directed toward developing *in vitro*^6^ and *in vivo*^7^ models to uncover promising therapeutic targets for ALS. Yet, the transition to effective therapies remains challenging. Stem cell-based approaches, especially those involving neural stem cells (NSCs), have shown great promise in delaying disease progression and reducing neuroinflammation^8^. NSCs have the unique ability to integrate into the central nervous system (CNS) parenchyma, form functional neuronal connections, and provide neuroprotective, anti-inflammatory paracrine effects^8,9^. Several pre-clinical and clinical studies have yielded encouraging results^10^, highlighting the potential of these cells in ameliorating ALS hallmarks and promoting transient positive results towards reduced neuroinflammation and improved motor function. However, key hurdles, including optimal cell source and dosage, transplantation timing, injection site and immunosuppressive regimen remain unclarified aspects^11^. Thus, non-clinical studies remain paramount to understanding ALS pathophysiology and uncovering the therapeutic potential of transplanted cells.

In a previous study^12^, we demonstrated that intraspinal transplantation of clinical-grade human-NSCs (hNSCs) in a rat model of ALS (SOD1^G93A^) can target multiple disease mechanisms, resulting in significant MN preservation, reduced neuroinflammation and extended animal survival^12^. Among others, these preclinical data have paved the way for translating this approach into clinical settings. Several phase I studies have successfully demonstrated positive, albeit transient, effects of hNSC transplantation in ALS patients^13–15^. Nonetheless, the limited size of enrolled patient cohorts often prevents from drawing definitive conclusions.

Subsequent phase II cell-dose-escalation studies have been conducted using the same strategy^11,15,16^. However, significant safety concerns and physical limitations have emerged. Specifically, the number of SC injections needed to increase the total transplanted hNSC count is constrained by the risk of backbone instability following surgery^11,16^. To address these limitations, we opted to implement an intracerebroventricular (ICV) transplantation strategy previously adopted for secondary progressive multiple sclerosis (SPMS) patients^17^. This approach involves transplanting hNSCs into the lateral ventricles via a standardized and less invasive surgical procedure. Importantly, this strategy may facilitate the delivery of a higher number of cells and a broader distribution throughout the neuraxis. Therefore, this study aims to outline challenges and promises of optimizing a non-clinical protocol to evaluate both safety and efficacy of hNSC ICV delivery as a novel and potentially more effective treatment for ALS. Here, we investigated the effects of unilateral and bilateral ICV transplantations using hNSCs at two different dosages: 3×10^5^ and 1×10^6^ cells. Both transient (15 days) and prolonged immunosuppression regimens, administered at low and high doses, were employed.

## Methods

### Ethical approval

All the experimental protocols involving live animals were performed in strict accordance with institutional guidelines in compliance with national (D.L. N.26, 04/03/2014) and international law and policies (new directive 2010/63/EU). The studies on nude and SOD1^G93A^ mice were approved by the Italian Ministry of Health (#751/2019-PR and #391/2021-PR protocols, respectively). Additionally, the ad hoc Institutional Animal Welfare Committee (OPBA) of the University of Milano-Bicocca and the Ethical Committee of the University of Turin approved these studies.

### Cell culture

hNSCs were isolated, expanded and characterized using the Neurosphere Assay as previously described^18,19^. Cells were seeded at a density of 10^4^ cells/cm^2^ in preconditioned medium in 25 (Corning, #CLS430639) or 75 cm^2^ flasks (Corning, #CLS430641U) (6 or 12 ml culture media, respectively), maintained in an incubator at 37 °C with >90% humidity, 5% O_2_ and 5% CO_2_ and allowed to proliferate as free-floating clusters named neurospheres. Neurospheres were mechanically dissociated through a p200 pipette to obtain the cell suspension that was replated at the same initial density. This step was routinely repeated and 24/48 h prior to transplantation in our animal models, cells were seeded at a density of 10^4^ cells/cm^2^. On transplantation day, hNSCs were harvested by centrifugation, and viable cells were counted by Trypan blue exclusion criteria. The appropriate cell number was then re-suspended in Hanks’ Balanced Salt Solution (HBSS, Carlo Erba, #FA30WL0607500).

### Animals

Two different animal models have been used in this study. Adult nude mice (Hsd:Athymic Nude-Foxn1^nu^, Envigo, #408761) have been employed to evaluate hNSC safety and biodistribution. The maintenance of the animal colony and behavioural assessments were conducted at the Department of Biotechnology and Biosciences, University of Milano-Bicocca, Italy. For efficacy studies, adult SOD1^G93A^ transgenic mice (B6SJL-Tg(SOD1*G93A)1Gur/J; The Jackson Laboratory, #002726) have been used. The maintenance of the animal colony and behavioural tests were conducted at the Neuroscience Institute Cavalieri Ottolenghi (N.I.C.O.), Department of Neuroscience “Rita Levi Montalcini”, University of Turin, Italy. Hemizygous transgenic progeny was obtained and maintained by crossbreeding SOD1^G93A^ transgenic males with wild type females (B6SJLF/1). Female mice were previously produced by crossing a C57BL/6J (B6) female mouse and an SJL/J (SJL) male mouse. Genotyping was performed by Polymerase Chain Reaction (PCR) on DNA extracted from mouse tails at approximately 21 days of age and by incubating a 0.5 cm long specimen of tail in 100 μL of lysis buffer (10 mmol/L Tris HCl, 50 mmol/L KCl, 0.01% gelatin, 0.45% IGEPALCA-630 (Sigma-Aldrich), 0.4% Tween-20) and 25 mg of proteinase K at 55°C overnight under gentle shaking. PCR allowed to detect the presence of *hSOD1* gene. The used primers were: 5’-CAT CAG CCC TAA TCC ATC TGA-3’, Transgene Forward, and 5’-CGC GAC TAA CAA TCA AAG TGA-3’, Transgene Reverse, for the hSOD1 gene and 5’-CTAGGC CAC AGA ATT GAA AGA TCT-3’and 5’-GTA GGT GGA AAT TCT AGC ATC ATC C-3’ for mouse interleukin 2 gene (mIL-2), as, respectively, Internal Positive Control Forward and Internal Positive Control Reverse. All animals were maintained in a virus and antigen-free facility with controlled temperature (18-22 °C), humidity (40-60 %), and 12-hour light-dark cycle. Food and water were provided *ad libitum*. All efforts were made to use the fewest number of animals and to minimize their suffering.

### Intracerebroventricular (ICV) hNSC transplantation

According to the different experimental purposes, mice were transplanted unilaterally (3×10^5^ cells/3 µl) or bilaterally (2 injections each, 5×10^5^ cells/5 µl) into the lateral ventricles (anteroposterior: -0.1 mm; lateral: ± 0.8 mm; dorsoventral: -2 mm from Bregma) with either hNSCs or HBSS as control, using a 30-gauge needle (Hamilton syringe) at a rate of 0.5 µl/30 sec.

For safety and biodistribution analysis a total of n=3 adult (5-8 weeks old) nude mice were used. For efficacy studies, at post-natal day 70 (P70; 0 days post-transplantation, dpt), mice were randomized into two experimental groups, hNSCs-treated and HBSS-treated (control group), and cyclosporin A (CsA; Sandimmune, Novartis) was administered to all animals subcutaneously on daily basis at 15 or 30 mg/kg starting from one day prior transplantation. Animals were subdivided as follows: EXP1 (Fig. 1A): SOD1^G93A^ mice, n=5 (3×10^5^ cells) 15 mg/kg CsA for 15 days sacrificed at both 20 (n=2) and 40 (n=3) dpt; EXP2: (Fig. 1B-C) SOD1^G93A^ mice, n=12 (3×10^5^ cells) *vs* n=11 (HBSS), 30 mg/kg CsA for 15 days sacrificed at the ES; EXP3: (Fig. 2A-G) SOD1^G93A^ mice, n=6 (1×10^6^ cells) *vs* n=6 (HBSS), 30 mg/kg CsA for 15 days sacrificed at the ES; EXP 4: (Fig. 2D-G): SOD1^G93A^ mice, n=6 (1×10^6^ cells) *vs* n=6 (HBSS), 30 mg/kg CsA for 15 days sacrificed 40 dpt. EXP 5: (Fig.3A-L) SOD1^G93A^ mice, n=13 (1×10^6^ cells) *vs* n=11 (HBSS), 30 mg/kg CsA for 40 days. Finally, post-transplantation cell viability and differentiation profile were qualitatively evaluated in a total of n=2 (7 dpt) and n=2 (15 dpt) SOD1^G93A^ mice (1×10^6^ cells), 30 mg/kg CsA for 15 days.

**Figure 1.**
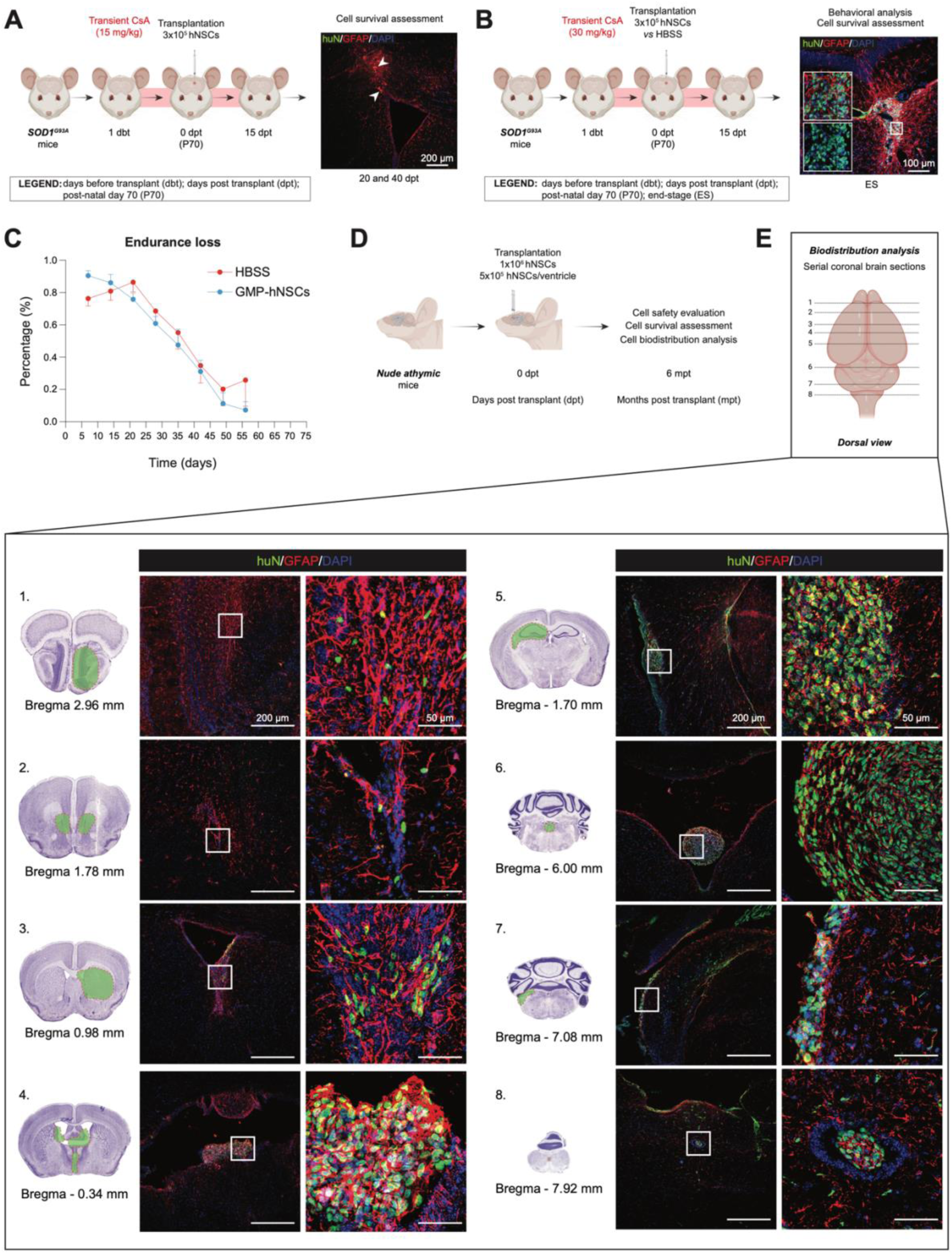
Preliminary evaluation of hNSC survival and efficacy under low-and high-dose transient immunosuppression, and long-term safety study of high-dose immortalized-hNSCs. (A) Experimental scheme showing unilateral hNSC transplantation (3×10^5^) timeline into the lateral ventricle of n=5 ALS mice (EXP 1) sacrificed at 20 (n=2) and 40 (n=3) dpt under low-dose (15 mg/kg daily) transient (15 days) immunosuppression (left) and representative confocal image of mouse brain showing only cellular debris (right) | huN, green; GFAP, red; DAPI, blue. (B) Experimental scheme showing unilateral hNSC transplantation (3×10^5^) timeline into the lateral ventricle of n=12 ALS mice under high-dose (30 mg/kg daily) transient (15 days) immunosuppression (left) (EXP 2) and representative confocal images of mouse brain showing huN^+^ living cells long-term (right) | huN, green; GFAP, red; DAPI, blue. (C) Endurance loss visualization of motor performance decline considering a total of n=12 mice + 3×10^5^ hNSCs (blue line) *vs* n=11 mice + HBSS (red line) (EXP 2). Data are expressed as mean values ± SEM. (D) Schematic showing the timeline of a bilateral ICV transplantation of 1×10^6^ immortalized-hNSCs in n=3 immunodeficient mice. (E) Upper panel: schematic representation of the different brain areas in which huN^+^ viable cells have been identified. | Lower panel: representative coronal brain sections from Paxinos et al., 2001 and confocal images of surviving huN^+^ (green) grafts 6 mpt. | huN, green; GFAP, red; DAPI, blue.

**Figure 2.**
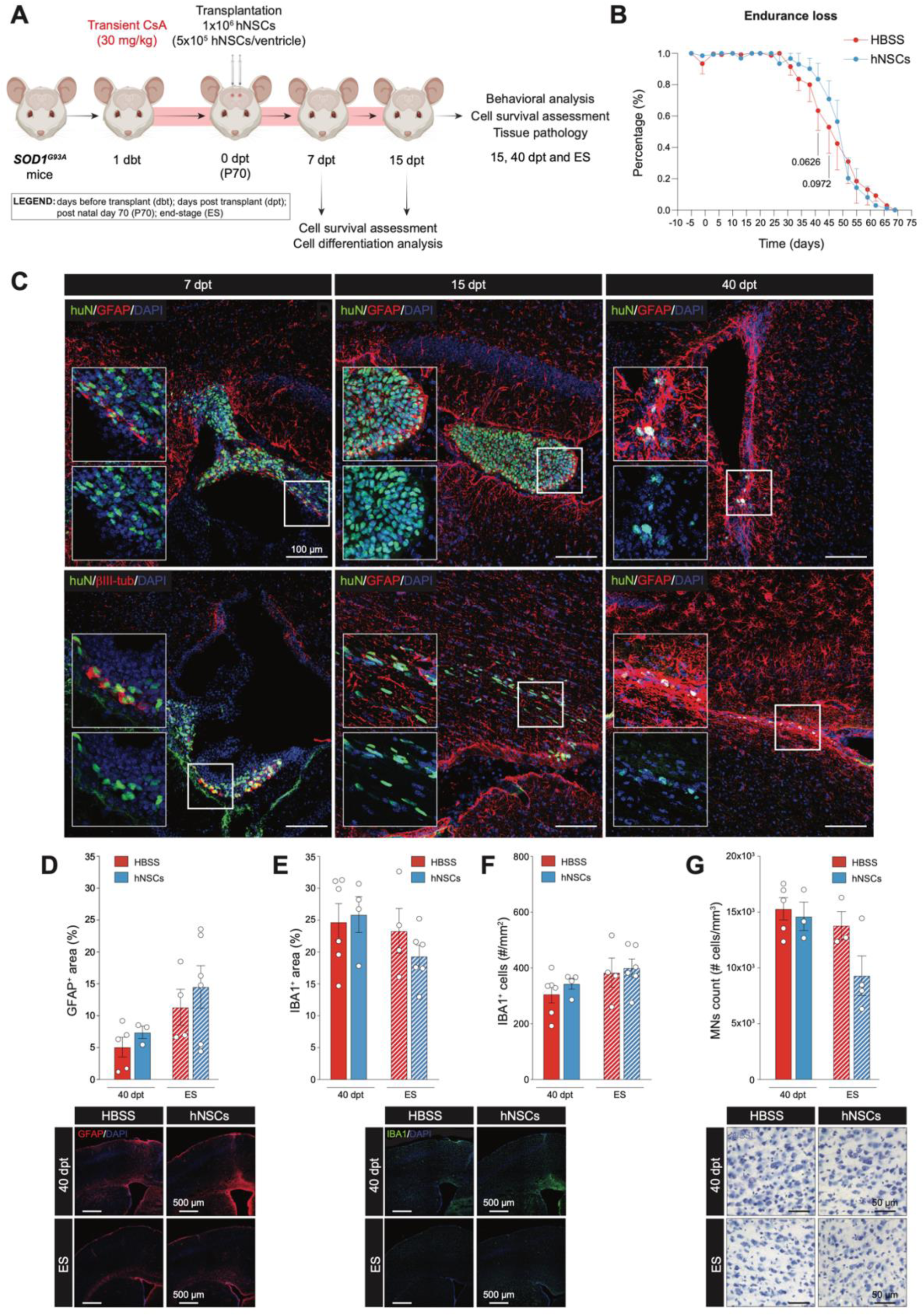
hNSC survival probability assessment and exploratory efficacy analysis of a bilateral ICV transplantation of high-dose hNSCs under high-dose transient immunosuppression. (A) Schematic showing the experimental setup of a bilateral high-dose (1×10^6^) hNSC ICV transplantation under high-dose (30 mg/kg daily) transient (15 days) immunosuppression regimen. (B) Endurance loss visualization of motor performance decline considering a total of n=6 SOD1^G93A^ mice + hNSCs (blue line) *vs* n=6 SOD1^G93A^ mice + HBSS (red line) (EXP 3); uncorrected Fisher’s LSD test, Mixed-effect model. Data are expressed as mean values ± SEM. (C) Representative confocal images showing the time-course assessment of post-transplantation hNSCs viability under a high-dose transient immunosuppression regimen. | huN, green; GFAP, red; DAPI, blue. (D) Neuropathological analysis of the M1-V layer’s astrogliosis at both 40 dpt and ES on hNSC- *vs* HBSS-treated mice; n≥3/time point/group (EXP 3 and 4) | GFAP, red; DAPI, blue. Data are expressed as mean values ± SEM. (E) Neuropathological analysis of the M1-V layer’s %Area of IBA1^+^ signal at both 40 dpt and ES on hNSC- *vs* HBSS-treated mice; n≥3/time point/group (EXP 3 and 4) | IBA1, green; DAPI, blue. Data are expressed as mean values ± SEM. (F) Neuropathological analysis of the M1-V layer’s assessed counting the total number of microglial cells per mm^2^ on hNSC- *vs* HBSS-treated mice at both 40 dpt and ES; n≥3/time point/group (EXP 3 and 4) | IBA1, green; DAPI, blue. Data are expressed as mean values ± SEM. (G) Primary MN stereological count in hNSC- *vs* HBSS-treated mice at 40 dpt and ES; n≥3/time point/group (EXP 3 and 4). Data are expressed as mean values ± SEM.

**Figure 3.**
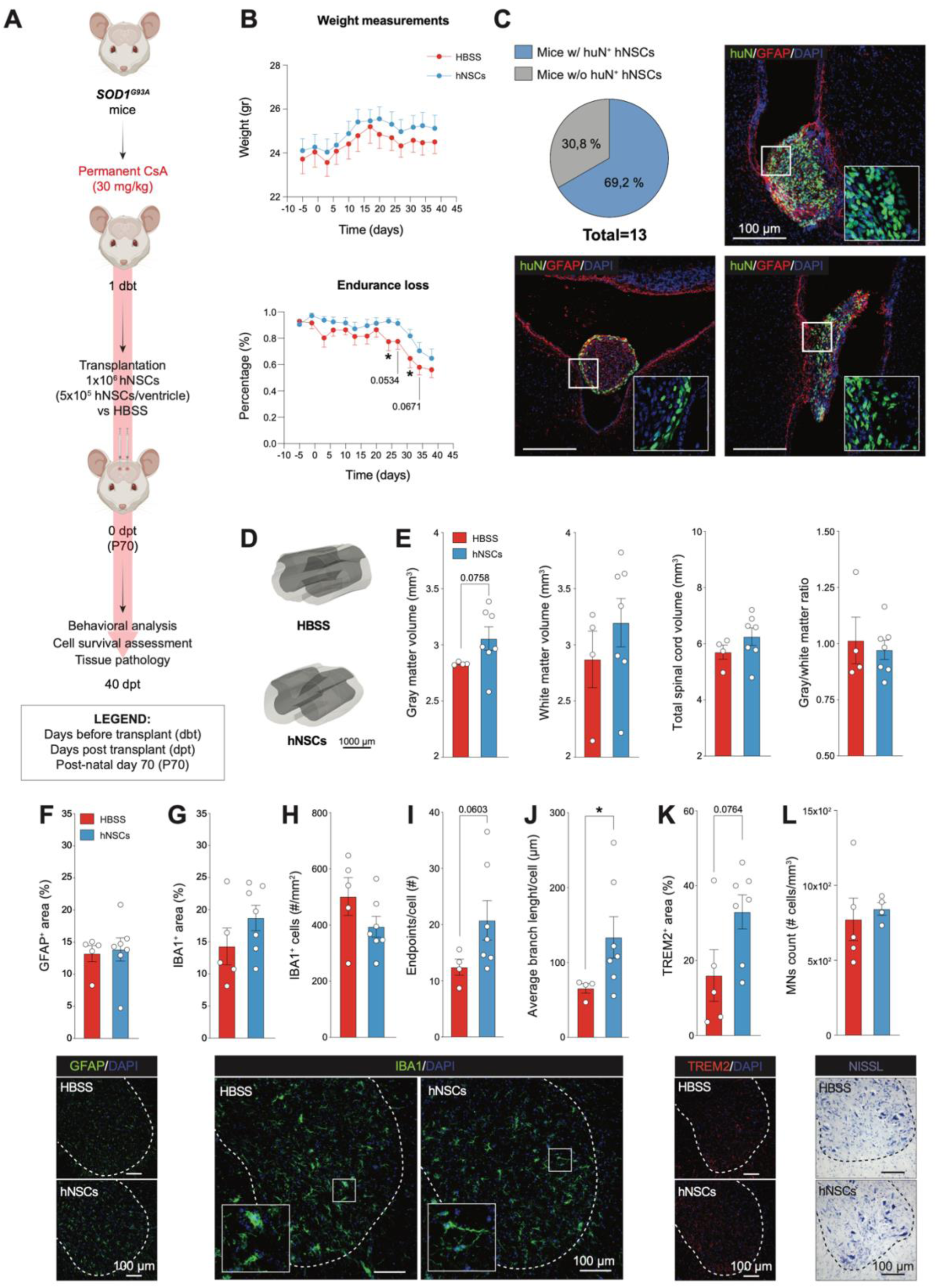
hNSC survival probability assessment and exploratory efficacy analysis of a bilateral ICV transplantation of high-dose hNSCs under high-dose prolonged immunosuppression. (A) Schematic showing the experimental setup of a high-dose hNSC bilateral ICV transplantation under high-dose prolonged immunosuppression regimen. (B) Weight measurements (upper graph) and endurance loss visualization of motor performance decline (lower graph) of hNSC- (n=13, blue line) *vs* HBSS-treated mice (n=11, red line) (EXP 5); uncorrected Fisher’s LSD test, Mixed-effect model (p-value: *≤0.05). Data are expressed as mean values ± SEM. (C) Pie chart showing the percentage of animals retrieved with and without huN^+^ living cells, and representative confocal images of huN^+^ living cells at an intermediate time point (40 dpt) upon high-dose prolonged immunosuppression regimen. | huN, green; GFAP, red; DAPI, blue. (D) Representative 3D reconstructions showing the analysed cervical SC of hNSC- *vs* HBSS-treated mice (n≥4/group; EXP 5). (E) Analysis of gray and white matter volume loss in the cervical tract of the SC of hNSC- *vs* HBSS-treated mice (n≥4/group; EXP 5). Data are expressed as mean values ± SEM. (F) Neuroinflammation analysis (astrogliosis) of the cervical tract of the SC of hNSC-and HBSS-treated mice at 40 dpt (n≥4/group; EXP 5) | GFAP, green; DAPI, blue. Data are expressed as mean values ± SEM. (G) Neuropathological analysis of the cervical tract of the SC assessed measuring microglial distribution in mice transplanted with hNSCs *vs* HBSS at 40 dpt (n≥4/group; EXP 5) | IBA1, green; DAPI, blue. Data are expressed as mean values ± SEM. (H) Analysis of the total amount of microglial cells (number of IBA1^+^ cells per mm^2^) in the ventral horn gray matter of the cervical SC of hNSCs- *vs* HBSS-transplanted mice at 40 dpt (n≥4/group; EXP 5) | IBA1, green; DAPI, blue. (I) *In vivo* morphological analysis of IBA1^+^ cells at 40 dpt assessed by total number of endpoints/cell; unpaired t-test with Welch’s correction (n≥4/group; EXP 5). Data are expressed as mean values ± SEM. (J) *In vivo* quantification of microglial branching at 40 dpt (n≥4/group; EXP 5); unpaired t-test with Mann-Whitney’s correction (p-value: *≤0.05). Data are expressed as mean values ± SEM. (K) Assessment of TREM2 protein levels in the cervical tract of the SC of hNSC- and HBSS-treated animals (n≥4/group; EXP 5); unpaired t-test with Welch’s correction. Data are expressed as mean values ± SEM. (L) MN stereological count in the cervical tract of the SC. hNSC- *vs* HBSS-treated mice at 40 dpt (n≥4/group; EXP 5). Data are expressed as mean values ± SEM

### Behavioural and disease progression assessment

Following hNSC grafting, all transplanted animals were monitored daily and, depending on experimental needs, tested weekly or twice a week until sacrifice. A clinical checklist was adopted to detect any behavioural and neurological sign of brain damage and/or tumor formation. Key parameters included significant weight loss (>20% respect to start of the treatment), skull deformation (hydrocephalus), exophthalmos, ataxia, and backbone curvature (kyphosis), assessed according to Guyenet et al. protocol^20^. To evaluate (i) disease progression; (ii) motor functions; and (iii) the potential efficacy of hNSC transplantation, rotarod performance was assessed, along with weight measurements, on transplanted and control SOD1^G93A^ mice starting from one week following transplantation or at P50 as described in the following paragraphs.

### Motor performance evaluation

The rotarod test was used to evaluate general motor coordination, strength and balance^21^. We first conducted a pilot experiment (EXP 1) during which animals were placed on the rotating rod at a speed starting from a minimum of 5 r.p.m. to a maximum of 32 r.p.m. being tested once a week, starting from one-week post-surgery. In all the following behavioural experiments (EXP 2, 3, and 5), mice were first trained for 20 days prior transplantation. Thus, animals were placed on the rotating rod at same speed described above starting from P50 and tested twice a week. Three days post-surgery, behavioural tests continued until sacrifice day. Each session consisted of three runs, with a maximum arbitrary cut-off time of 300 seconds per run. The best performance time per test was used for further analysis. For statistical analysis, endurance loss was measured by dividing each mouse rotarod score at different time points by its highest recorded score.

### Weight measurements

Each mouse was weighed (in grams) once or twice a week with a standard scale until sacrifice. For subsequent analysis, the mean weight of all mice in each experimental group was calculated.

### Animal sacrifice procedure and tissue processing

To collect both brain and spinal cord (SC), animals were sacrificed by overdose of anaesthesia and perfused with 4% paraformaldehyde (PFA; Sigma #158127) in a phosphate buffer (PBS 1X, Sigma-Aldrich, #D8537) solution at different time points accordingly to experimental needs: 7, 15, 20, 40 dpt, and ES. Tissues were then removed, placed 2 h in 4% PFA at 4 °C, cryopreserved in sucrose gradients (10% for 24 h, 20% for 24 h and 30% until the samples were fully imbibed), embedded in Optimal Cutting Temperature (OCT; Killik, Bio Optica, #05-9801) medium and frozen above liquid nitrogen vapours. To perform histological analysis, tissues were serially sectioned on a cryostat.

For the hNSC safety and biodistribution analysis in nude mice, serial coronal sections of the brain and brainstem, and transverse spinal SC sections were cut at the thickness of 20 µm and directly placed on the surface of glass slides (epredia, #J1800AMNZ). For SOD1^G93A^ samples, a free-floating cutting method was employed. Serial brain (coronal) and SC (transverse) sections were cut at the thickness of 30 µm and collected in a cryopreserving solution for further analysis.

### Immunofluorescence and immunohistochemistry

#### Immunofluorescence on glass slides

Coronal brain and transverse SC sections were placed on glass slides (epredia) and immunohistochemistry was performed for cell survival, differentiation and biodistribution analysis on nude mice. Blocking Solution (PBS-1X with 10% Normal Goat Serum (NGS) and 1% TRITON X-100 was added at room temperature on each slide. Sections were then incubated in a humid chamber overnight at 4 °C with the appropriate primary antibodies diluted in serum solution (NGS 10% and PBS 1X), respectively: anti-human Nuclei (huN) (1:200, Ms, IgG1, Millipore, MAB 1281), anti-Glial Fibrillary Acidic Protein (GFAP) (1:500, Rb, IgG, Dako Z0334) and anti-βIII Tubulin (1:400, Rb, IgG, BioLegend, 802001). The day after, tissues were washed with PBS 1X and then, incubated for 90 minutes at room temperature with the appropriate secondary antibodies (Abs II) in serum solution, as described before. The following Abs II were used: Alexa Fluor 488 (1:1000, α-Ms, Jackson, #115-545-205) and Alexa Fluor 594 (1:1000, α-Rb, Invitrogen, #A11012). Cell nuclei were labelled with 4’,6- Diamidin-2-phenylindol (DAPI, 1:1000, Roche Diagnostics GmbH, Mannheim, Germany, #10236276001) in PBS 1X for 20 min at room temperature and the slides mounted using coverslips with aqueous mounting (Fluor Save Reagent, Calbiochem, #CALB345789-20).

#### Immunofluorescence on free floating sections

Neuroinflammation analysis on SOD1^G93A^ mice was performed via immunofluorescence following the free-floating protocol. Samples were rinsed 3 times with PBS 1X on tilting plate. A permeabilization step followed with the addition of 0.3% TRITON X-100 and PBS 1X for 30 min at room temperature on tilting plate. Blocking solution (10% NGS, 0.3% TRITON X-100, PBS 1X) was added for 30 minutes at room temperature on tilting plate. Then, sections were incubated overnight at 4 °C with anti-GFAP (1:500, Rb, IgG1, Dako, Z0334), anti-IBA1 (1:500, Rb, IgG, Wako, 019-19741), and anti-TREM2 (1:100, Rat, IgG2b, Biotechne R&B System, MAB 17291) antibodies. The next day, after washing with PBS 1X, the following Abs II were used: Alexa Fluor 594 (1:500, α-Rat, Invitrogen, A11007), and Alexa Fluor 488 (1:1000, α-Rb, Jackson, 111-545-144). Cell nuclei were labelled with 4’,6-Deamidine-2’-phenylindole dihydrochloride (DAPI, 1:1000; Roche, #10236276001) for 20 min at room temperature. After washing with PBS 1X, samples were mounted on glass slides (epredia), left to dry and then gently washed with deionized water (dH2O). Slides were cover-slipped with aqueous mounting as before.

#### Nissl staining

Brain and SC sections were Nissl-stained to perform motor neurons (MNs) stereological count. Respectively, serial coronal brain or transverse SC sections were hydrated in distilled water for 1 min before staining in 0.1% Cresyl Violet acetate for 10 min, then dehydrated in an ascending series of ethanol (EtOH 95% and EtOH 100%) for 30 sec, cleared in xylene for 30 sec for 2 times and cover-slipped with Eukitt (Bioptica, Milan, Italy, #W01030706). Sections were mounted on 4% gelatine-coated Superfrost slides (epredia) and air-dried overnight at room temperature.

#### hNSC safety, survival, differentiation, and biodistribution analysis

The safety, survival, differentiation, and biodistribution of hNSCs transplanted into the lateral ventricles of nude mice were evaluated qualitatively by serial Z-stack confocal images of different part of the brain, brainstem and SC. Nikon A1R confocal microscope was used for image acquisition.

#### Motor cortex neuroinflammation analysis

All the images of serial coronal sections throughout the rostro-caudal extent of the M1 motor cortex-V layer were obtained by performing large-image acquisitions with Nikon A1R confocal microscope using the 10x objective and processed with Software NIS Elements A1 Version 5.30.06. The lambda (λ) parameter (optical configuration) was set on three channels as follows: DAPI (HV=90, Offset=-80, Laser=4.0), FITC (HV=60, Offset=-80, Laser=3.5) and TRITC (HV=40, Offset=-70, Laser=2.0). In order to obtain an image containing all the motor cortex (M1), the large image parameter was set to be 2×2 or 3×3, depending on ROI (Region Of Interest) dimension. Z-stack of 28μm thickness with 10 steps sampling were acquired from the top to the bottom of each section, with an acquisition interval of 3.11 μm each. Consecutive Z-stack images were converted to a maximum intensity projection (MIP) picture and the final Z-stack confocal image was obtained for further examinations.

Quantitative neuroinflammation analysis was conducted on brain samples of both HBSS- and hNSCs-treated SOD1^G93A^ mice at different time points: 0 (P70), 40 dpt, and ES. Z-stack confocal images of coronal sections in TIFF format were analysed with Fiji ImageJ (ImageJ2, Version 2.9.0/1.53t) and Photoshop (23.5.0 Release). Firstly, by employing ImageJ software, the region of interest (ROI) of M1-V layer was manually drawn on confocal images based on both DAPI staining and the additional guide of *The Mouse Brain Atlas in Stereotaxic Coordinates*^22^.

For each channel, the % of Area (%Area) of the GFAP^+^ and IBA1^+^ signal was quantified on thresholded and non-thresholded images. For thresholded images, the %Area is the percentage of pixels in the ROI that has been highlighted in red by using the “*Mean*” Threshold parameter. For non-thresholded images, the %Area is the percentage of non-zero pixels. For our analysis, only the %Area of thresholded images was considered. For statistical analysis, a minimum of 3 and a maximum of 14 serial coronal sections/animal were analysed and the plot value was calculated as follows: (sum of all the areas for the thresholded images / sum of all the areas for the non-thresholded images)*100.

Additionally, the number of microglial (DAPI^+^/IBA1^+^) cells per mm^2^ was quantified and included in our analysis. A manual, non-stereological count was performed on the previously described confocal images by mean of Count tool on Photoshop or ImageJ. To normalize the obtained data, the final plot value/animal was obtained as follows: (sum of the counted cells in each section / total non-thresholded ROIs area in mm^2^).

#### SC neuroinflammation analysis

Confocal images of transverse sections of the cervical (C1-C8) tract of the SC were acquired at 20x objective using a Nikon A1R confocal microscope. Four sections per cervical segment of each animal were selected for quantification analysis and the plot value was calculated as follows: (sum of all the areas for the thresholded images / sum of all the areas for the non-thresholded images)*100. Identical camera and fluorescent light source settings were used across all sections and for each paired antibodies. The lambda (λ) parameter (optical configuration) was set on three channels as follows: DAPI (HV=70, Offset=-50, Laser=5.0), FITC (HV=50, Offset=-80, Laser=3.5) and TRITC (HV=32, Offset=-95, Laser=4.0) for IBA1/TREM2 antibodies; and DAPI (HV=70, Offset=-50, Laser=5.0) and FITC (HV=30, Offset=-90, Laser=3) for GFAP antibodies. Z-stack of 15μm thickness with 8 steps sampling were acquired from the top to the bottom of each section, with an acquisition interval of 2.14 µm each. Consecutive Z-stack images were converted to a maximum intensity projection (MIP) picture and the final Z-stack confocal image was obtained for further examinations.

Using ImageJ software, the %Area of the GFAP^+^, IBA1^+^, and TREM2^+^ signal was measured within a constant ROI (size: 487.01×408.13 µm) centred in the grey matter of the ventral horn. The quantification was performed on thresholded and non-thresholded images. Moreover, the number of microglial (DAPI^+^ /IBA1^+^) cells per mm^2^ was quantified following the same approach as per the motor cortex analysis. Finally, a morphological assessment of microglial cells (number of endpoints/cell and average branch length/cell in µm) was included in our analysis employing a validated protocol for 20x immunofluorescence images^23^. Briefly, skeleton analysis was performed using Fiji ImageJ v.2.14.0. software plugins AnalyzeSkeleton (2D/3D) after applying the following criteria: pixel radius = 3 and mask weight = 0.6, and pixel radius = 0.5 and threshold = 50 for outliers removal. Finally, the cut-off value of 2.583 was identified for further analysis. Data are expressed as the average number of endpoints per cell, and average process length in μm per cell from n≥4 ROIs (1 per each SC section) per mouse from n≥4 independent biological replicates (±SEM).

#### Motor neuron stereological counting

Upper and lower motor neuron (MN) quantification was performed on both hNSC- and HBSS-treated SOD1^G93A^ mice at different time points, 40 dpt and ES. The motor cortex (M1’s layer V) and the ventral horns of cervical (C3-C5) spinal segments have been evaluated. One series of Nissl-stained serial brain and SC sections (one every 180 µm) from each group was analysed. The estimation of MN density was obtained by using a stereological technique, the Optical Fractionator, with a computer-assisted microscope and the Stereo-Investigator software (MicroBrightField, Williston, VT, USA). Cells were counted on the computer screen with the use of an Optronics MicroFire digital camera mounted on a Nikon Eclipse E600 microscope. MNs’ nucleoli were counted at 40x. Only neurons with a diameter ≥ 10 µm (classified as large pyramidal cells in the cerebral cortex) and ≥ 16 µm (considered alpha MNs in the SC) and located in the appropriate position were counted. The total volume of the reconstructed segment (expressed in mm³) and the MN density (expressed as number of MNs/mm^3^) were calculated Additionally, grey and white matter volumes were evaluated by Neurolucida software (Version 9) on four serial transverse sections (1 every 720 µm) sampled within the cervical tract.

#### Statistical analysis

Statistical analysis was conducted using GraphPad Prism Software (Version 9.5.0, GraphPad Software, La Jolla, CA, USA). Normal distribution of individual values was verified prior to each test, and outliers were excluded following an outliers identifier test (ROUT method, Q=5%). For comparisons between 2 groups and >1 condition, a Two-Way ANOVA (mixed-effect model) followed by Fisher’s LSD test was performed. One-way ANOVA (Brown-Forsythe and Welch ANOVA tests) were applied for comparisons of >2 groups and 1 condition. For comparisons between 2 groups under 1 condition, a parametric unpaired t-test with Welch’s correction or a nonparametric unpaired t-test with Mann-Whitney’s correction was used. All data are presented as mean ± SEM (Standard Error of the Mean), and significance was determined at *p* ≤ 0.05 (*).

## Results

### Unilateral ICV injection of low-dose hNSCs under high-dose transient immunosuppression shows suboptimal cell survival in SOD1^G93A^ mice

Previous data by our group demonstrated the safety and differentiation profile of extensively characterized^24^ GMP-hNSCs upon unilateral ICV transplantation in immune-deficient adult mice^17^.

Starting from these results, we investigated the survival and integration of hNSC line in immunocompetent SOD1^G93A^ mice under a mild-dose (15 mg/kg daily) transient (15 days) immunosuppression regimen with CsA. We transplanted a total of 3×10^5^ cells unilaterally at approximately 70 days of age (P70) (EXP 1), a stage when histological markers of the disease are already present, though motor symptoms are still negligible^25–27^. When sacrificed at 20 and 40 dpt we did not observe surviving cells via immunostaining for the human specific antibody (huN). This indicates that a mild immunosuppression was insufficient for hNSC survival in mice (**Fig. 1A**). To improve hNSC engraftment, we employed the highest immunosuppression dosage recommended for mice (30 mg/kg daily)^28^ for 15 days upon injection of the same hNSCs dose (**Fig. 1B, left**). Motor functions and weight were monitored until End Stage (ES). In this second experiment (EXP2), huN^+^ hNSCs were detected in 3 out of 12 ES mice, mostly near the ventricles (**Fig. 1B, right**). However, we did not observe any behavioural improvement in motor performances (**Fig. 1C**), weight decline or survival rate in transplanted SOD1^G93A^ mice *vs* controls (SOD1^G93A^ mice + HBSS). Altogether, these results suggest that, although increasing immunosuppression enhanced hNSC survival, the cell dosage was still suboptimal to contrast disease progression.

### Bilateral ICV transplantation of high-dose immortalized hNSCs is feasible and safe

We then investigated whether increasing the dosage of transplanted hNSCs could ameliorate histological and behavioural hallmarks in ALS mice. Here exploited the greater *in vitro* amplification potential, as well as promising therapeutic abilities^29,30^ of an immortalized hNSC line previously characterized in our lab^19^. We first assessed the safety of transplanting a high dose of hNSCs through a long-term study in nude athymic mice (**Fig. 1D and E**). A total of 1×10^6^ immortalized hNSCs were injected into the lateral ventricles of adult (5-8 weeks) nude mice via bilateral ICV injection (5×10^5^ cells/ventricle). Animals were euthanized at 6 mpt (**Fig.1D**), and gross pathological analysis showed no signs of inflammatory or neoplastic processes such as alterations of the cerebral parenchyma and ventricles asymmetry. Moreover, serial brain, brainstem and SC sections, allowed us to identify a robust migratory capacity of transplanted hNSCs, reaching the 3^rd^ and 4^th^ ventricles up to the central canal of the cervical SC. Of note, hNSCs were detected adhering to the ventricular wall and migrating into the parenchyma, reaching multiple brain structures such as olfactory bulbs, striatum, lateral septum, corpus callosum and CA1 region of the hippocampus. The most rostral and caudal CNS regions in which we identified transplanted hNSCs were Bregma 2,96 mm and Bregma -7,92 mm, respectively (based on *The Mouse Brain Atlas’*^22^ annotations, **Fig.1E**). Overall, our results demonstrate that transplantation of a high dose (1×10^6^) immortalized hNSCs via bilateral ICV injection in immunodeficient hosts is feasible and safe.

### Bilateral ICV transplantation of high-dose hNSCs under high-dose transient immunosuppression shows trend in delayed motor decline

To investigate the potential benefits of high-dose hNSC transplantation, we next transplanted SOD1^G93A^ mice bilaterally with 1×10^6^ hNSCs at P70 under a high-dose (30 mg/kg daily) transient (15 days) immunosuppression regimen (**Fig. 2A**). We first assessed motor functions and animal survival until ES and observed a slight delay in motor decline in the hNSC-treated group *vs* controls, particularly between 41 (0.84±0.10% *vs* 0.64±0.13%, respectively; p=0.0626) and 45 dpt (0.71±0.12% *vs* 0.53±0.17%, respectively; p=0.0972) (**Fig. 2B**; EXP 3). We then performed a qualitative time-course analysis of post-transplantation cell viability and differentiation profile (**Fig. 2C**). At 7 dpt, we observed both huN^+^/βIII-tub^+^ and huN^+^/GFAP^+^ hNSCs, indicating their *in vivo* differentiation into neuronal and astrocytic lineages. Moreover, viable huN^+^ hNSCs persisted up to 15 dpt and migrated into the brain parenchyma showing a “stretched” morphology (**Fig. 2C**). However, at later time points (40 dpt and ES), we observed viable hNSCs inside the third ventricle of only one ES mouse (data not shown). When huN^+^ cells were not detected, cellular debris were identified near the injection site and within the cerebral parenchyma (**Fig. 2C**), indicating suboptimal long-term survival of hNSCs under transient high-dose immunosuppression.

Despite the low long-term survival rate of hNSCs, we investigated whether their persistence into the host CNS was sufficient to induce motor cortex histopathological changes. We first assessed both immunofluorescence GFAP^+^ and IBA1^+^ signals, indicative of astro-and micro-gliosis, respectively, in the V-layer of the murine primary motor cortex (M1) and found no major differences both at 40 dpt and ES (**Fig. 2 D-F**). These findings suggest that the level of hNSC persistence achieved with transient high-dose immunosuppression was insufficient to induce substantial histopathological changes in the motor cortex. Moreover, at 40 dpt and in both groups (hNSCs *vs* HBSS; EXP 4), we observed a marked increase in the %GFAP^+^ area in some brain sections, predominantly near the injection site, but occasionally extending rostrally into the murine cortex (**Fig. 2D, lower 40 dpt panels**). This increase likely resulted from the surgery-induced focal astrogliosis, potentially confounding and limiting any cortical neuropathological analysis at intermediate time points. By ES, despite the overall GFAP^+^ signal was higher (**Fig. 2D**), this effect was reduced, even becoming undetectable in some mice, suggesting it was transient and spatially confined (**Fig. 2D, lower ES panels**). Next, we evaluated the MN degeneration rate in the same anatomical region via stereological count and obtained comparable number of MNs between hNSC-treated and control groups at both 40 dpt and ES (**Fig. 2 G**). These results indicate that high-dose (30 mg/kg) transient (15 days) immunosuppression did not ensure optimal cell survival following high-dose (1×10^6^) hNSC transplantation. However, we observed a promising trend of transient behavioural improvement at around 40 dpt, despite persistent astro-and microgliosis and ongoing primary MN degeneration. Overall, these findings suggest that hNSC transplantation has potential for delaying motor impairments, although further optimization of cell survival is necessary to induce concomitant histopathological changes and achieve more sustained therapeutic outcomes.

### Bilateral ICV transplantation of high-dose hNSCs under high-dose prolonged immunosuppression delays motor symptoms degeneration

We next tested whether an increased dose (30 mg/kg daily) and extended duration (until sacrifice) of immunosuppression would enhance behavioural and histopathological outcomes in SOD1^G93A^ mice following ICV transplantation of 1×10^6^ hNSCs (EXP 5; **Fig. 3A**). Focusing on animals sacrificed at 40 dpt, an intermediate time point where prior studies^12^ suggest the greatest pathological amelioration, and consistent with results previously shown in Fig. 2B, we observed significantly improved motor function in the hNSC-treated group *vs* controls starting from 24 dpt (0.93±0.03% *vs* 0.78±0.07%, respectively) and lasting until 31 dpt (0.82±0.05% *vs* 0.65±0.07%, respectively) without affecting weights in both groups (**Fig. 3B**). Lastly, we detected huN^+^ hNSCs in 9/13 (∼70%) transplanted animals, extending to the Silvius aqueduct and 4^th^ ventricle (**Fig. 3C**). These findings suggest that extended immunosuppression guarantee optimal hNSC survival, contributing to significant motor improvements.

We then assessed whether hNSCs transplanted under prolonged immunosuppression induced histopathological changes. Here, we focused our analysis on the cervical and lumbar SC tracts, both described as particularly vulnerable regions to neurodegeneration in ALS pathology^31,32^. Importantly, this approach also avoids the confounding surgery effects seen in our earlier motor cortex analysis. We first measured gray (3.06±0.10 mm^3^, hNSCs *vs* 2.83±0.006 mm^3^, HBSS; p=0.0758) and white matter (3.20±0.21 mm^3^, hNSCs *vs* 2.87±0.25 mm^3^, HBSS) volumes of SOD1^G93A^ mice and found a trend toward reduction in cervical SC degeneration in hNSC- transplanted mice *vs* controls (**Fig. 3D-E**). While the total cervical SC volume in the hNSC group showed a non-significant trend indicative of lower parenchymal degeneration *vs* controls (6.25±0.31 *vs* 5.70±0.25 mm^3^, respectively), the gray-to-white matter ratio remained consistent across groups, potentially suggesting comparable disease effect in both areas (**Fig. 3D-E**). Overall, these findings might suggest a potential for hNSC-induced neuroprotection in mitigating cervical SC degeneration as indicated by trends in gray and white matter volume preservation.

To investigate a potential neuroprotective mechanism exerted by transplanted hNSCs, we assessed astro-and micro-gliosis levels in the ventral horn gray matter of the same SC tract. By looking at the GFAP^+^ astrocytes, we observed both groups to have similar astrocyte coverage suggesting that transplanting high-dose (1×10^6^) hNSCs under a prolonged high-dose immunosuppression did not impact astrogliosis (**Fig. 3F**). To investigate whether hNSC transplantation affect microgliosis, we assessed IBA1^+^ cells distribution (**Fig. 3G**), number (**Fig. 3H**) and morphology (**Fig. 3I-J**) in the hNSC-treated *vs* control group. Interestingly, we observed a trend toward increasing IBA1^+^ %Area (18.73±1.97% *vs* 14.30±2.87%, respectively; **Fig. 3G**) alongside a decrease trend in the total number of IBA1^+^ cells (394.00±37.24 cells/mm^2^ vs 501.60±67.16 cells/mm^2^, respectively; **Fig. 3H**). This suggests that while microglial cell number is reduced, they occupy a larger area within the ventral horn gray matter, potentially reflecting reduced microgliosis and a shift to a more ramified, anti-inflammatory, phenotype. Consistently, IBA1^+^ cell morphological analysis revealed a higher number of endpoints per cell (20.70±3.50 *vs* 12.40±1.44; p=0.0603; **Fig. 3I**) and significantly longer branch lengths (133.60±27.62 *vs* 65.50±6.26 µm; **Fig. 3J**) in the hNSC *vs* HBSS group, respectively. This is indicative of less activated microglial morphology suggesting reduced cervical SC inflammatory environment in hNSC-transplanted ALS mice. To further explore this hypothesis, we assessed TREM2^+^ microglia levels within the same anatomical region (**Fig. 3K**). Here, we show a trend toward increased %TREM2^+^ area compared to the HBSS control group (33.03±4.5% *vs* 16.03±6.8%, respectively; p=0.0764; **Fig. 3K**). This may support TREM2 crucial role in promoting a neuroprotective microglia phenotype. Finally, we examined whether the observed microglial changes were coupled with a higher MN preservation in the cervical SC (**Fig. 3L**). Alpha MN count showed a non-significant trend towards reduced neurodegeneration in the hNSC group *vs* controls (879.70±31.97 vs 773.40±141.80 cells/mm^3^, respectively), thus further corroborating potential hNSC-mediated neuroprotective effects. Of note, no differences were observed in the lumbar SC (data not shown), confining positive results to anatomical regions directly reached by transplanted cells.

Collectively, these findings indicate that high-dose (1×10^6^) hNSC transplantation in SOD1^G93A^ mice under sustained high-dose (30 mg/kg) immunosuppression mitigates endurance loss. Furthermore, despite being injected distally in the lateral ventricles, hNSCs show potential to reduce microglia activation in the cervical SC, highlighting their widespread anti-inflammatory effects.

## Discussion

Currently, no therapy can reverse or halt ALS progression. Given the severity of the prognosis, there is an urgent need to translate promising approaches into effective clinical treatments. Here, we aimed to optimize key experimental parameters to establish an alternative, non-clinical ICV transplantation protocol of hNSCs for ALS. Our findings demonstrate that under a prolonged, high-dose immunosuppression hNSCs survive, spread along the neuraxis, and occasionally migrate into the cerebral parenchyma of SOD1^G93A^ mice. Furthermore, transplanted hNSCs delays motor symptoms degeneration and reduce spinal cord microgliosis in this ALS mouse model. To our knowledge, this is the first study to show that foetal-derived hNSCs transplanted into the lateral ventricles may induce behavioural and histopathological changes ALS. Moreover, our work reveals the low immune tolerance of mice to transplanted hNSCs, suggesting that alternative models, such as rats, may provide a more suitable environment for long-term integration studies of human cells^12^.

Previous intraspinal hNSC transplantations in SOD1^G93A^ models showed beneficial effects, including partial protection of spinal MNs, delayed motor function degeneration, and extended survival^12,33–37^. Although clinical studies have partially confirmed these results, the benefits proved to be transient, likely due in part to the aggressive nature of ALS and the underestimation of optimal cell dosage^13,14^. In this context, increasing the number of transplanted cells via multiple intraspinal injections in SOD1 mice has been reported to cumulatively enhance therapeutic effects^35^, thus reinforcing the rationale that suboptimal dosing may limit hNSC efficacy. However, translating this strategy to clinical practice is hindered by the risks associated with the extensive laminectomy and invasiveness required for multiple SC injections in humans.

In this study, we hypothesized that ICV transplantation could offer a novel, less invasive alternative, leveraging CSF circulation to enhance hNSC biodistribution and neurotrophic factors release throughout the neuraxis. Additionally, it would allow for repeated injections, ensuring continuous delivery of healthy hNSCs, potentially maximizing their therapeutic impact. Of note, this procedure has already been standardized for intracerebral drug delivery in humans and was recently applied in our clinical study, where up to 24 million cells were safely administered into the ventricles of SPMS patients^17^.

Here, we report that only higher doses of hNSCs (1×10^6^), combined with prolonged high-dose (30 mg/kg) immunosuppression, enhanced cell survival, biodistribution, and therapeutic outcomes in ALS mice. Interestingly, cells spread along the ventricles, improving from the low-to-high dose treatment, and reached the central canal of the cervical SC, though none migrated near the thoraco-lumbar tract or into the spinal parenchyma. While the astro-and micro-gliosis evaluation in the motor cortex was confounded by the gliotic scar-like reaction to the needle, we observed a trend toward higher spinal volume and reduced microgliosis in the cervical region, identified by a more ramified microglial morphology in the hNSC-treated mice. Under disease conditions, microglia adopt a pro-inflammatory phenotype, further exacerbating neurodegeneration^38^. Although recent studies highlight the dynamic and heterogenous nature of microglial cells, challenging the traditional resting (M2) and activated (M1) phenotypes, it remains clear that microglia undergo pronounced morphological changes from a ramified to an ameboid shape in CNS pathologies^38^ which seems to be counteracted following hNSC transplantation. This evidence was coupled with higher TREM2 levels, a crucial regulator of microglial phagocytic activity with recognized anti-inflammatory properties^39^, in the same SC tract. Elevated levels of this protein have been implicated to a not yet fully clarified neuroprotective mechanisms in the inflamed milieu. Both *in vitro*^40^ and *in vivo*^41^ studies have demonstrated decreased TREM2 expression across several neurodegenerative diseases models, including ALS^42^. However, the limited understanding of TREM2’s function across ALS stages prevents drawing definitive conclusions. Overall, these behavioural and histopathological changes suggest that the local presence of viable cells, rather than direct parenchymal integration, may be sufficient for therapeutic benefits (at least to a certain extent). This aligns with previous studies on mesenchymal stem cell transplantation into the cisterna magna^43,44^ or lumbaris^45^ of ALS mice, reinforcing the potential of paracrine effects over direct cell replacement. Further studies should explore whether multiple ICV injections or subsequent injections in different CNS regions (ICV and intrathecal, e.g. cisterna lumbaris) would be more effective compared to a single administration. Finally, the observed limited parenchymal migration may reflect insufficient inflammatory signals required for NSC homing, which may not yet be present at biologically relevant concentrations during early disease stages when we delivered our hNSCs^46^. Nonetheless, our decision to transplant pre-symptomatically was twofold: (i) evidence suggests that upper MNs may undergo pathology prior to clinical symptoms onset, making them early targets in ALS^25–27,47,48^; and (ii) early transplantation maximizes the time window for hNSCs within the hosts environment, accounting for their slower inner biological features respect to their murine counterparts and ALS progression in mice. Furthermore, it is known that neurodegeneration and functional decline in ALS patients can precede diagnosis^49^. Thus, early transplantations might extend hNSC therapeutic window and potentially slow down the degeneration of upper MNs. However, investigating the effects of ICV cell delivery at later stages, when inflammation is more robust, may enhance hNSC homing and neuroprotection and be more clinically relevant as ALS diagnosis in humans typically occurs at symptomatic stages.

Inevitably, there are limitations to our study, primarily related to the reactive murine immune response to exogenous human cells. This immune reactivity hampers our ability to precisely determine the extent and timing of hNSC death, as well as assess whether immune responses may have compromised their therapeutic efficacy. Additionally, the slow intrinsic biology of human cells in relation to the rapid disease progression of our model, along with suboptimal immunosuppression, further limits the potential for a sustained therapeutic effect^36,50^.

In conclusion, our findings support the potential of hNSCs to mitigate ALS pathology. However, optimizing the delivery method, selecting the correct animal model, and targeting specific tissues is essential to maximize the benefits of cell-based therapies. Although SOD1^G93A^ mice provide a well-established model for ALS research, SOD1 mutations account for only a small fraction of human ALS cases, and many therapies that showed promise in this model have failed in clinical trials^51^. Moving forward, comprehensive investigations involving both *in vitro* and *in vivo* models, encompassing different ALS mutations and sporadic cases, will be crucial to elucidate the molecular mechanisms underlying hNSC therapeutic effects.

## Author contribution

Conception and design of the study: DF, MB, and MG. Acquisition of data: IL, EV, CF, DPC, DMR, SdlM, RMC, EP, SL, and EL. Analysis and interpretation of data: IL, EV, CF, DF, MG, and MB. Drafting or revising the manuscript: IL, EV, CF, DF, DMR, MG, MB, SM, JR, and AV.

